# Basketball interest as gateway to STEM interest: Testing a large-scale intervention to enhance STEM interest in sports-engaged populations

**DOI:** 10.1101/2025.03.14.643324

**Authors:** Emily J. Hangen, Amy K. Loya, John F. Drazan

**Affiliations:** SUNY Brockport; Union College; Fairfield University

**Keywords:** STEM education, Informal learning, Interest, Work-force development, Sports Science, Motivation

## Abstract

**Background:** To address documented demographic disparities in Science, Technology, Engineering, and Math (STEM) disciplines, targeted interventions have been designed to close achievement gaps and remedy the “leaky STEM pipeline”. However, much less work has focused on designing complementary interventions that to seek to broaden the population of youth who “enter the pipeline” or who are exposed to STEM and interest is stoked from the onset. The current work seeks to broaden STEM engagement in youth by developing a STEM intervention using unrecognized forms of cultural capital. Rather than situating STEM learning within traditional “STEM-Intensive” programs that require a pre-existing level of STEM interest, the current work developed and tested interventions designed to broaden engagement of youth with pre-existing interest in sports. These basketball-based STEM education interventions utilized sports as a venue for informal STEM learning. The current work tested 3 variations of this novel STEM education intervention: a single hour event, a one-day clinic, and a multi-day camp. Each of these programs was delivered to naturally-occurring, targeted populations of students with high levels of pre-existing sport engagement and low levels of STEM engagement. The sample populations included students from 2nd grade to 12th grade, with middle school students being the most heavily represented.

**Results:** Interest in pursuing a STEM career significantly increased across all 3 variations of the basketball-based STEM education intervention. Notably, these effect sizes were descriptively larger for programs of longer duration (e.g. multi-day camp vs. one hour event) and were significantly stronger for students who reported playing basketball prior to the program.

**Conclusions:** The current work introduces a novel STEM education intervention that utilizes students’ pre-existing interest (e.g. basketball) as a bridge to STEM engagement. This approach was shown to be repeatedly successful in enhancing participants’ STEM career interest, especially for participants from underserved communities with high levels of sport interest.

## Introduction

Lack of diversity within the STEM career fields has negative implications for the rate of technical innovation (Campbell et al., 2013; Østergaard et al., 2011; Valantine et al., 2016) and for distribution of economic opportunities for youth (Melguizo & Wolniak, 2012; Rothwell, 2013). The process of STEM training is often described in metaphor as the “STEM Pipeline”, which consists of a mix of formal and informal STEM learning experiences. Formal STEM learning takes place in the school environment and is shown to influence persistence in STEM disciplines. For example, rigorous precollege formal STEM preparation appears to help to retain students in collegiate STEM programs (Moses et al., 2011); however, a personal interest and passion for STEM, as demonstrated by voluntarily taking additional STEM classes in high school, regardless of rigor, is a stronger predictor of four-year STEM degree completion than specific enrollment in rigorous advanced placement coursework (Sadler et al., 2014). Informal STEM experiences take place out of formal school settings, such as robotics clubs or visits to science museums (Allen & Peterman, 2019; Eshach, 2007; Subotnik et al., 2009). Informal STEM engagement is also beneficial for increasing student engagement, scholastic achievement, and ownership of STEM topics (Bell et al., 2009; Gottfried & Williams, 2013). In addition, youth who participate in informal STEM programs, such as robotics clubs, are significantly more likely to pursue STEM majors in college (Dabney et al., 2012; Eguchi, 2016; Miller et al., 2018). Taken together, the most important role that pre-collegiate STEM experiences (both formal and informal) play in the production of future STEM practioners appears to be developing and retaining a student’s personal interest and identity in STEM rather than imbuing advanced technical competency (Maltese et al., 2014; Maltese & Tai, 2011).

Key processes consistent across STEM disciplines are self-directed inquiry and problem-solving, yet the majority of opportunities for informal STEM engagement are content-focused, “STEM-Intensive” programming such as robotics clubs, science fairs, visiting science museums, or coding summer programs. The content-focused nature of these programs can dissuade participation of youth who do not have pre-existing interest in the content. Evidence shows that youth who are already interested in STEM are more likely to voluntarily spend their available time in such programs, which leads to self-selecting participation where STEM-predisposed youths are more likely to engage in these valuable programs (Dabney et al., 2012; Miller et al., 2018). In fact, researchers have documented that a notable challenge of evaluating informal STEM education programs is the “ceiling effect” among participants who have such high scores entering the program that their scores cannot be meaningfully improved (Staus et al., 2021). These researchers have noted how the ceiling effect is particularly common in informal STEM education programs:

> *“This effect is often attributed to the biased nature of participation. Informal science learning opportunities, including after school programs, are particularly susceptible to this effect due to the fact that participants generally choose to participate because they are already interested in and potentially knowledgeable about the content area*.*”* (p. 2, Staus et al., 2021).

As noted, there is a fundamental limitation with the present STEM pipeline—informal STEM learning opportunities are typically reliant on voluntary participation, which is highly influenced by pre-existing interest and awareness on the part of the learner. Critically, this means that informal STEM learning opportunities are unlikely to reach those in most need of an authentic introduction to STEM: youth who are not already aware of and interested in STEM fields. As such, the present STEM pipeline appears to be designed to accelerate and retain STEM-predisposed youths into STEM careers rather than broadening engagement among youth who may not be initially drawn to these “STEM-Intensive” programs.

STEM-Intensive programs often require existing interest, parental support, and prior STEM knowledge to participate or excel, all of which are influenced by cultural capital. Cultural capital, first proposed by Pierre Bourdieu (1973), represents the skills, knowledge, and experiences that are valued within a given field and its institutions and may offer clues to these patterns. In STEM education, cultural capital often comes in the form of exposure to hands-on technical activities like coding, robotics, or lab-based science early in life, familial role models and support for STEM, and strong self-confidence in technical tasks (Fletcher Jr. & Carroll, 2024; Read et al., 2019). Cultural capital also reflects the values of gatekeepers and dominant stakeholders within a given field and shapes who has access to opportunity and who thrives within a given field (Stolle-McAllister, 2011; Tilbrook & Shifrer, 2022). Archer and colleagues introduced the concept of “Science Capital” to capture science-related forms of social and cultural capital and developed a survey to quantify the distribution of scientific capital across 3, 658 secondary school students in England (Archer et al., 2015). They found that scientific capital is disproportionately accessible to students with higher generalized cultural capital (parental high school completion and university attendance, visits to museums, and number of books in the home). 50% of the students with the highest measurement of science capital (5% of the entire sample) reported their desire to study science in college while only 6% of the students with the lowest amount of science capital (27% of the sample) reported college science aspirations. Without the appropriate forms of cultural and social capital, many youths can view the existing STEM-Intensive programs as inaccessible or culturally-foreign, a gap that traditional STEM programs frequently fail to address. As such, the forms of cultural capital that are fungible within the existing STEM outreach programing implicitly rewards students who already possess STEM cultural capital, reinforcing existing underrepresentation and maintaining the status quo.

The current work seeks to address this mismatch of cultural capital to enhance the desirability of participating in informal STEM programming for the broader population of youth who may lack exposure to STEM programming that fits their interests. Previously researchers have sought to identify different types of unrecognized cultural capital that are endemic to these underserved populations and make them fungible within informal STEM programming (Eglash & Bennett, 2009). Developing programs that make these forms of presently unrecognized capital fungible within informal STEM is an important, but underutilized, approach to broaden participation. One approach is the development of Culturally Situated Design tools focused on identifying authentic embodiments of STEM within community cultures and practices such as algorithmic procedures in cornrow hairstyling, coordinate systems in Navajo loom use, and even hip-hop rhythmic patterns (Babbitt et al., 2015; Emdin et al., 2016; Eglash et al., 2006; Eglash & Bennett, 2009; Eglash et al., 2013).

The current work builds upon past work which has centered on basketball, and sports more broadly, as an attractive source of unrecognized capital for informal STEM engagement (Drazan et al., 2014, 2015, 2016). Sports provide an area of shared interest wherein STEM professionals and youth can deploy STEM analysis techniques to learn about something deeply meaningful to youth athletes: how to improve as an athlete. Athlete identity and passion for sports can serve as a robust and enduring entryway into STEM where athletes already possess a growth mindset (McCarthy et al., 2008). For more details on the benefits of sports for STEM education, please see (Drazan, 2020). Our past work has shown that the application of STEM techniques such as data visualization and analytics to basketball shooting heatmaps results in higher levels of STEM interest, sports-science interest, and a higher confidence within their own sport (Drazan et al., 2017). We have also shown that in a short sport science skills assessment showcase, also known as a combine, youth athletes experienced statistically significant increases in the same indices through the provision of custom sports performance training advice based on individual combine data collection results (Loya et al., 2023a). This study included both students who participated in pre-existing informal STEM program and youth athletes, both of whom participated in the sport science combine. Interestingly, we observed the same “ceiling effect” of STEM interest among the already STEM engaged youth as prior informal STEM education programs (Staus et al., 2021).

Importantly, a primary outcome we will examine is how offering youth a culturally-situated informal STEM program will affect youth’s interest in pursuing STEM disciplines. The focus on interest as an outcome is deliberate because Expectancy-Value theory (Eccles, 2009) suggests that making material personally-relevant (which is the aim of providing a culturally-situated STEM program) can enhance student motivation and be an effective way to enhance students’ participation and success in STEM. According to Expectancy-Value theory (Eccles, 2009), students’ motivation and academic choices are the product of their expectations for success and their subjective value of the task. Among these subjective values is utility value, or the perceived usefulness of the task to the individual (Eccles, 2009). As empirical support for the application of this theory, utility value interventions have been shown to successfully increase the likelihood of college students persisting in a STEM major (Canning et al., 2018) and enhance students’ semester GPAs and grades in an undergraduate biology course, especially among first-generation, underrepresented minority students (Harackiewicz et al, 2014).

Additionally, interest is a primary outcome in this study because educational psychology has placed particular focus on the construct of interest and has found that individuals who repeatedly experience situational interest, or whose learning context temporarily sparks interest, can develop more long-lasting individual interest in that topic (Rotgans & Schmidt, 2017). Because this culturally-situated exposure to STEM is designed to highlight the personal relevance of STEM by tying it to a pre-existing interest (e.g. in basketball), we anticipate that it will elicit situational interest in STEM which has the potential for developing into longer-lasting individual interest in STEM. Lastly, interest is an important outcome to examine since it is a critical construct for examining how learning transfers across contexts (Pugh & Bergin, 2006), which can be especially relevant when translating knowledge acquired between informal STEM programs and formal STEM education.

We were approached to develop a large-scale implementation of our programs in the Midwest by SC Johnson. Together with 4^th^ Family, our long-term non-profit community partner, we created the SC Johnson and 4^th^ Family STEM and Sport Team Up Initiative that was delivered in the spring and summer of 2022. As program directors, we built upon our previous work in sport biomechanics for STEM education to implement a novel, large scale STEM education intervention. The program was assessed for efficacy using an anonymous, matched pre-posted questionnaire with all participants. The study procedures across this and all studies in this current work were reviewed and were granted an IRB exemption from the Human Subjects Review Committee at Union College as per 45 CFR 46.104(d)(2).

The purpose of the current study is to evaluate the efficacy of sports for informal STEM education across three separate informal STEM programs of increasing duration (1 hour, 4 hours, and 3 days) across a large population of youth in different community contexts. We hypothesized that the basketball focus of this STEM program would preferentially engage youth with a pre-existing interest and identity within basketball. Further, we hypothesized that STEM interest increases would increase with respect to the length of the program.

## Study 1

The Science of the Slam was designed to engage students in scientific methods and expose them to the personal usefulness and relevance of STEM by connecting STEM practices to a prevalent pre-existing student interest in basketball. Specifically, unlike traditional STEM extracurriculars (e.g. robotics clubs, science fairs, etc.), the Science of the Slam was intentionally designed to engage basketball-interested youth in STEM by helping students recognize the overlap between processes in STEM and basketball. Importantly, this program was developed by the authors who have joint expertise as scholars in STEM fields and in basketball as two out of three authors were collegiate basketball players and remain active in the basketball community.

To investigate intervention efficacy, we examined changes in students’ STEM motivation. Specifically, we assessed interest in pursuing a STEM career, enjoyment of STEM classes, and self-efficacy in STEM. We also investigated whether students’ self-efficacy and interest in sports would be affected by the Science of the Slam.

## Methods

### Participants

Over the course of 12 days a total of 2675 students recruited from five middle schools in Racine, Wisconsin and four middle schools on the south side of Chicago participated in the Science of the Slam (*M*_*Age*_ = 12.63 years, *SD*_*Age*_ = 0.99 years). 1986 students completed the pre-survey, 1546 students completed the post-survey, and 947 students completed both surveys. The majority of the students (96%) were in middle school (see Table 1).

**Table 1.** Grade Level Distribution.

| Grades | Study 1<br>Frequency | Study 2<br>Frequency | Study 3<br>Frequency |
| --- | --- | --- | --- |
| 2<sup>nd</sup> | 4 | 12 | -- |
| 3<sup>rd</sup> | -- | 7 | -- |
| 4<sup>th</sup> | -- | 29 | 13 |
| 5<sup>th</sup> | 71 | 27 | 21 |
| 6<sup>th</sup> | 835 | 18 | 22 |
| 7<sup>th</sup> | 868 | 18 | 13 |
| 8<sup>th</sup> | 855 | 32 | 16 |
| 9<sup>th</sup> | -- | 7 | 8 |
| 10<sup>th</sup> | -- | 6 | 2 |
| 11<sup>th</sup> | -- | 3 | 2 |
| 12<sup>th</sup> | -- | 2 | -- |
| Not reported | 42 | -- | 1 |
| Total | 2675 | 161 | 98 |

Notably, only 7% of sample reported any current involvement in STEM extracurriculars. In contrast, the majority of the sample played sports, with 65% of the sample reporting that they played sports (see Table 2). In particular, basketball was the most popular sport that students played (28% of sample played basketball; See Table 3). As added evidence of pre-existing sports interest, 20% of the sample reported that they aspired to become a professional athlete (see Table 4).

**Table 2.** Participation in General Activities.

| Activity | Study 1 | Study 2 | Study 3 |
| --- | --- | --- | --- |
|  | Percentage (N) | Percentage (N) | Percentage (N) |
| Sports | 65% (1295) | 97% (127) | 93% (78) |
| Performing Arts | 43% (859) | 21% (28) | 25% (21) |
| STEM | 7% (138) | 11% (15) | 12% (10) |
| Other | 8% (168) | 8% (11) | 7% (6) |

**Table 3.** Participation in Specific Activities.

| Activity | Study 1 | Study 2 | Study 3 |
| --- | --- | --- | --- |
|  | Percentage (N) | Percentage (N) | Percentage (N) |
| Video Gaming | 35% (701) | 34% (45) | 26% (22) |
| Basketball | 28% (559) | 76% (99) | 87% (73) |
| Art or Design classes | 25% (490) | 14% (18) | 11% (9) |
| Football | 19% (383) | 35% (46) | 38% (32) |
| Band/Play an instrument | 16% (322) | 11% (14) | 13% (11) |
| Soccer | 16% (322) | 15% (20) | 15% (13) |
| Baseball or Softball | 13% (250) | 13% (17) | 23% (19) |
| Sing/Choir | 12% (247) | 2% (3) | 4% (3) |
| Track and Field | 12% (246) | 23% (30) | 20% (17) |
| Swimming/Diving | 12% (237) | 13% (17) | 11% (9) |
| Dance/Gymnastics | 10% (198) | 6% (8) | 1% (1) |
| Martial Arts | 6% (129) | 4% (5) | 4% (3) |
| Theatre/Improv | 6% (121) | 2% (2) | 6% (5) |
| Cheerleading/Majorette | 6% (116) | 4% (5) | 1% (1) |
| Tennis | 5% (92) | 3% (4) | 6% (5) |
| Coding Classes | 5% (90) | 5% (7) | 8% (7) |
| Wrestling | 4% (89) | 5% (6) | 8% (7) |
| Student Government | 4% (70) | 7% (9) | 2% (2) |
| Hockey or Ice Skating | 3% (57) | 4% (5) | 4% (3) |
| Lego League/Club | 3% (53) | 2% (3) | 2% (2) |
| Chess Club | 3% (51) | 3% (4) | 4% (3) |
| Robotics Club | 2% (46) | 3% (4) | 4% (3) |
| Science Fairs | 2% (38) | 2% (3) | 1% (1) |
| Debate Club | 1% (21) | 2% (3) | 0% (0) |
| Field Hockey | 1% (19) | 5% (7) | 1% (1) |

**Table 4.** Participants’ Career Aspirations.

| Career | Study 1 | Study 2 | Study 3 |
| --- | --- | --- | --- |
|  | Percentage (N) | Percentage (N) | Percentage (N) |
| eSports/Professional Gamer | 25% (499) | 26% (34) | 26% (22) |
| Video Game Designer/Tester | 24% (481) | 23% (30) | 15% (13) |
| Artist | 24% (467) | 18% (24) | 13% (11) |
| Actor | 23% (448) | 11% (15) | 19% (16) |
| Chef/Baker | 21% (410) | 11% (14) | 7% (6) |
| Professional Athlete | 20% (401) | 36% (47) | 54% (45) |
| Business Owner/Executive | 20% (391) | 17% (22) | 17% (14) |
| Doctor | 17% (340) | 9% (12) | 11% (9) |
| Graphic Designer | 16% (309) | 8% (11) | 11% (9) |
| Lawyer | 15% (307) | 10% (13) | 11% (9) |
| Influencer/Celebrity | 15% (289) | 9% (12) | 15% (13) |
| Engineer | 13% (267) | 13% (17) | 19% (16) |
| Musician/Singer | 13% (261) | 6% (8) | 10% (8) |
| Nurse | 13% (261) | 5% (6) | 4% (3) |
| Military | 12% (236) | 8% (11) | 7% (6) |
| Teacher/Professor | 12% (234) | 7% (9) | 8% (7) |
| Coach | 11% (219) | 26% (34) | 24% (20) |
| Police Officer/Detective | 11% (219) | 8% (11) | 5% (4) |
| Computer Programmer/IT | 10% (193) | 8% (10) | 10% (8) |
| Mechanic | 9% (186) | 5% (7) | 8% (7) |
| Writer/Author | 9% (180) | 6% (8) | 4% (3) |
| Scientist | 8% (167) | 11% (14) | 15% (13) |
| Veterinarian | 8% (158) | 4% (5) | 0% (0) |
| Ballerina/Dancer | 8% (152) | 3% (4) | 0% (0) |
| Realtor | 7% (141) | 2% (3) | 7% (6) |
| Architect | 5% (104) | 5% (7) | 6% (5) |
| Full-time Parent/Homemaker | 5% (103) | 1% (1) | 2% (2) |
| Pilot | 5% (97) | 5% (6) | 6% (5) |
| Construction Worker | 4% (89) | 4% (5) | 2% (2) |
| Astronaut | 4% (82) | 4% (5) | 4% (3) |
| Marketer | 4% (82) | 4% (5) | 2% (2) |
| Farmer | 3% (68) | 1% (1) | 0% (0) |
| Paleontologist | 2% (45) | 3% (4) | 0% (0) |
| Politician | 2% (31) | 2% (3) | 1% (1) |
| Biomechanist | 0% (0) | 0% (0) | 0% (0) |

### Materials

Students rated their agreement to items across all measures on a 1 - *Strongly Disagree* to 5 - *Strongly Agree* Likert scale.

#### STEM motivation variables

STEM career interest was computed as the average response to 2 items, “I am interested in studying STEM-related subjects in high school or college” and “I am interested in pursuing a job or career in a STEM-related field” for both pre-intervention STEM career interest (*α* = .73, McDonald’s *ω* = 0.74) and post-intervention STEM career interest (*ω* = .80, McDonald’s *ω* = 0.80). STEM enjoyment was computed as the average agreement rating to 2 items including “I enjoy math and/or science-related activities” for both pre-intervention STEM enjoyment (*ω* = .80, McDonald’s *ω* = 0.81) and post-intervention STEM enjoyment (*α* = .82, McDonald’s *ω* = 0.83). Self-efficacy in STEM was assessed with a single item “I am good at math and/or science” for both pre and post intervention assessments.

#### Sports science motivation variables

All sports science motivation measures were assessed with single items. Interest in sports science was assessed by student agreement ratings to the statement “I want to learn more about how I can use math and science to become a better athlete”. Value of sports science was assessed by agreement ratings to the statement, “I believe using math and science can make me a better athlete.”

#### Sport motivation variables

Sport enjoyment was computed as the average agreement rating to 2 items such as “I enjoy playing sports or participating in other athletic activities” for both pre-intervention sport enjoyment (*α* = .87, McDonald’s *ω* = 0.87) and post-intervention sport enjoyment (*α* = .90, McDonald’s *ω* = 0.89). Self-efficacy for sports was measured with a single item, “I am good at playing sports or other athletic activities”, for both pre and post intervention sports self-efficacy.

#### Participant extracurriculars and career aspirations

In addition to the measures previously described, students reported their regular participation in after-school activities and career aspirations.

### Procedure

Students completed all measures twice, prior to and after participating in the Science of the Slam. The Science of the Slam consisted of a 1-hour event where students were guided to use the scientific method to predict which contestant would win a Slam Dunk contest that was delivered across school gymnasiums to audiences of 50 – 150 students at a time. We recruited 3-5 local high school, junior college, and college basketball players who serve as “dunkers” for the activity. In brief, student participants were taught the scientific method through the use of slam dunk contest where the vertical jump height of each dunker was measured using DIY Arduino tools that were built by student volunteers that use the flight time method (Drazan et al., 2016). Students then generated a hypothesis that was two-fold: 1) the best dunker will jump the highest 2) the best jumper will be identified as the best dunker by eliciting the loudest cheer from the student audience.

We also focused on the importance of the testability and falsifiability of a given hypothesis. We did not know in advance which dunker would win the event, so in the course of the program students were exposed to how hypotheses are not always correct (i.e. the crowd’s favorite dunker may not be the participant who jumped the highest during vertical jump height testing). In this way, we introduce large groups of students to the scientific method, experimental design, biomechanics, and engineering while providing a rich, interactive experience for student participants. Following the dunk contest, students had the opportunity to measure their own vertical jump height to compare their results to the newly crowned slam dunk champion. The program concluded with a discussion of biomechanics and muscle physiology as a way to introduce students to how to improve their own testing metrics and imbue them with curiosity around both sport performance and STEM principles. This program was repeated each class period (8-10 times a day) to make the program accessible to every student within a given school.

### Data analysis

We tested for initial group differences in interests by running independent t-tests on data gathered before the start of the Science of the Slam program. Next, we tested for Science of the Slam effects by conducting paired t-tests (for composite scores) and Wilcoxon signed ranks tests (for single-item measures) on pre and post survey responses. Lastly, we ran mixed ANOVA’s to test whether program effects were moderated by participant characteristics. Specifically, we examined whether the impact of the program depended on 1) whether students reported already participating in STEM extracurriculars prior to the Science of the Slam, 2) whether students played sports, and 3) whether students participated in basketball in particular. For our analyses across all 3 studies, we used R (Version 4.3.3; R Core Team, 2024) and the R-packages *coin* (Version 1.4.3; Hothorn et al., 2006, 2008), *dplyr* (Version 1.1.4; Wickham, François, et al., 2023), *effsize* (Version 0.8.1; Torchiano, 2020), *knitr* (Version 1.46; Xie, 2015), *ltm* (Version 1.2.0; Rizopoulos, 2006), *MASS* (Version 7.3.60.0.1; Venables & Ripley, 2002), *MOTE* (Version 1.0.2; Buchanan et al., 2019), *msm* (Version 1.7.1; Jackson, 2011), *papaja* (Version 0.1.2; Aust & Barth, 2023), *polycor* (Version 0.8.1; Fox, 2022), *psych* (Version 2.4.3; Revelle, 2024), *scales* (Version 1.3.0; Wickham, Pedersen, et al., 2023), *survival* (Version 3.5.8; Therneau & Grambsch, 2000), and *tinylabels* (Version 0.2.4; Barth, 2023).

Attention check questions were included to ensure data quality (Maniaci & Rogge, 2014). students who gave the same agreement rating to the items “I enjoy math and/or science-related activities” and “I do not enjoy math and/or science-related activities” were removed from analysis (with the exception of students who answered “Neutral” to both). This resulted in the removal of 251 students for a new total of 2424 students.

## Results

### Initial group differences

As expected, students who were already participating in STEM extracurriculars had greater pre-intervention STEM career interest (*M* = 3.50, *SD* = 0.93) than their peers (*M* = 3.10, *SD* = 0.93), *t*(132.05) = -4.51, *p* < .001, *d* = 0.44 (see Figure 1).

**Figure 1.**
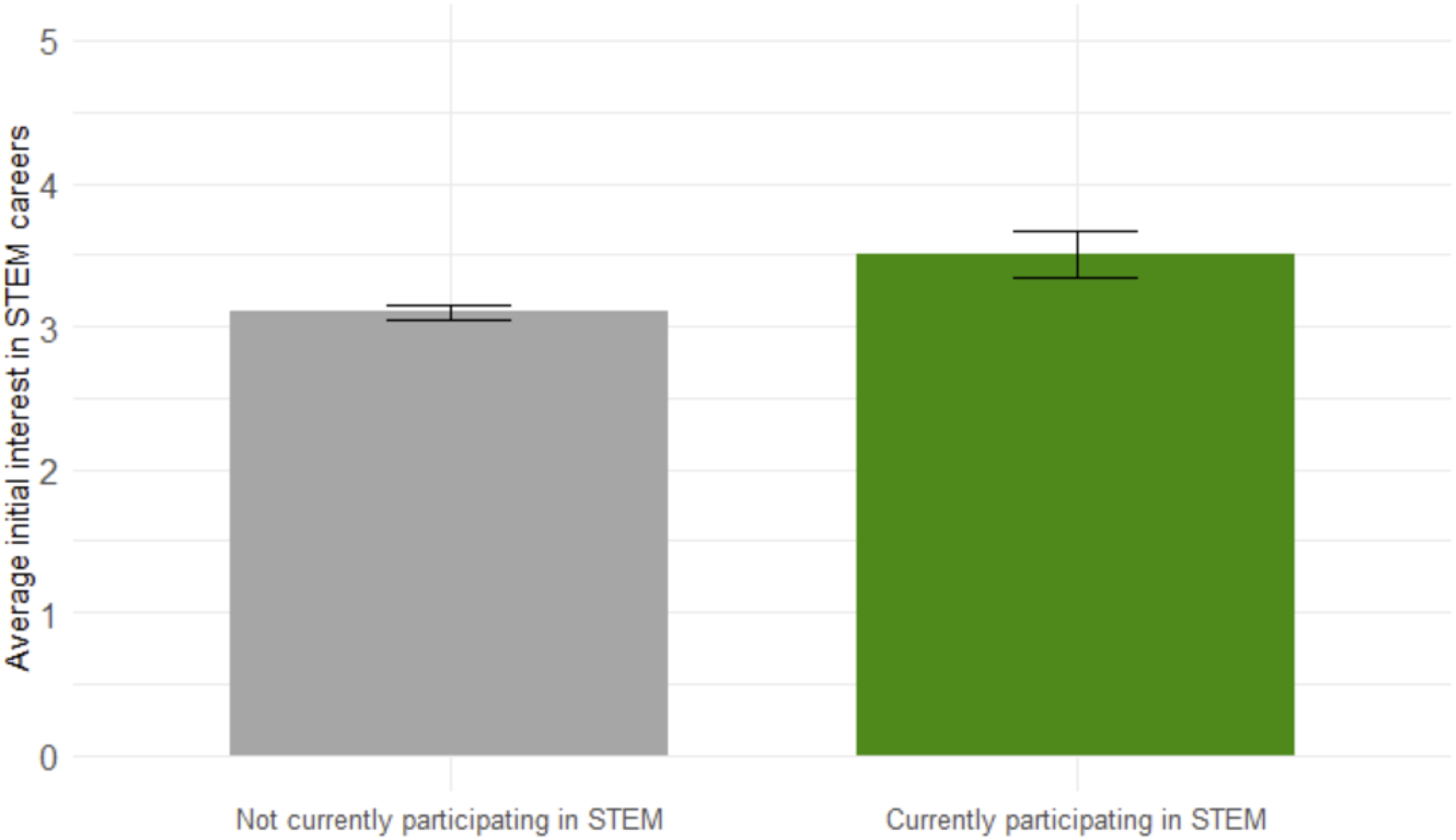
Initial Interest in STEM careers differs based on STEM activity. **Note:** Most participants were not currently participating in STEM (n = 1467) compared to students currently participating in STEM activities (n= 138).

### Effects on STEM Motivation

#### STEM career interest

There was a significant increase in STEM career interest from before participation in Science of the Slam (*M* = 3.12, *SD* = 0.94) to after (*M* = 3.25, *SD* = 0.97), *M*_*D*_ = 0.15, 95% CI, [0.09, 0.21], *t*(798) = 4.87, *p <* .001, *d* = 0.16 (see Figure 2). This increase in STEM career interest significantly differed between students who didn’t play sports and those who did, *F*(1.946.12) = 28.36, *p <* .001. STEM career interest did not significantly change for students who did not play sports, *t*(276) = −0.36, *p* = .721, however, STEM career interest significantly increased for students who did play sports, *M*_*D*_ = 0.25, 95% CI [0.16, 0.33], *t*(466) = 5.28, *p* < .001, *d* = 0.27.

**Figure 2.**
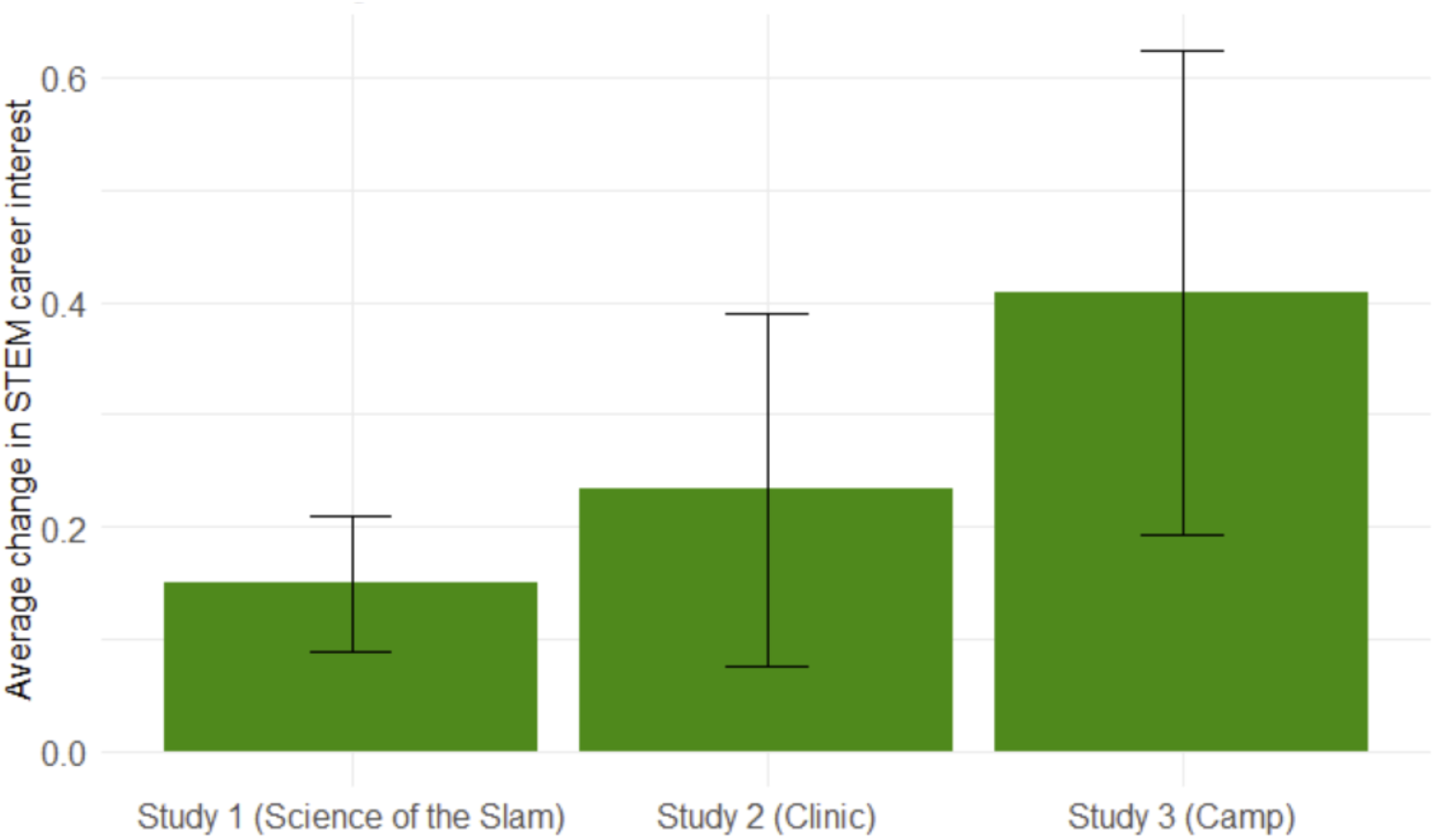
Average increase in STEM career interest across all 3 studies.

The increase in STEM career interest was also strongest among basketball players, *F*(1.975.28) = 18.59, *p* < .001. Students who did not play basketball showed a small but significant increase in STEM career interest, *M*_*D*_ = 0.08, 95% CI [0.02, 0.15], *t*(548) = 2.41, *p* = .0.16, *d* = 0.09, however, basketball players had a larger increase in STEM career interest, *M*_*D*_ = 0.33, 95% CI [0.190.47], *t*(194) = 4.72, *p* < .001, *d* = 0.36 (see Figure 3).

**Figure 3.**
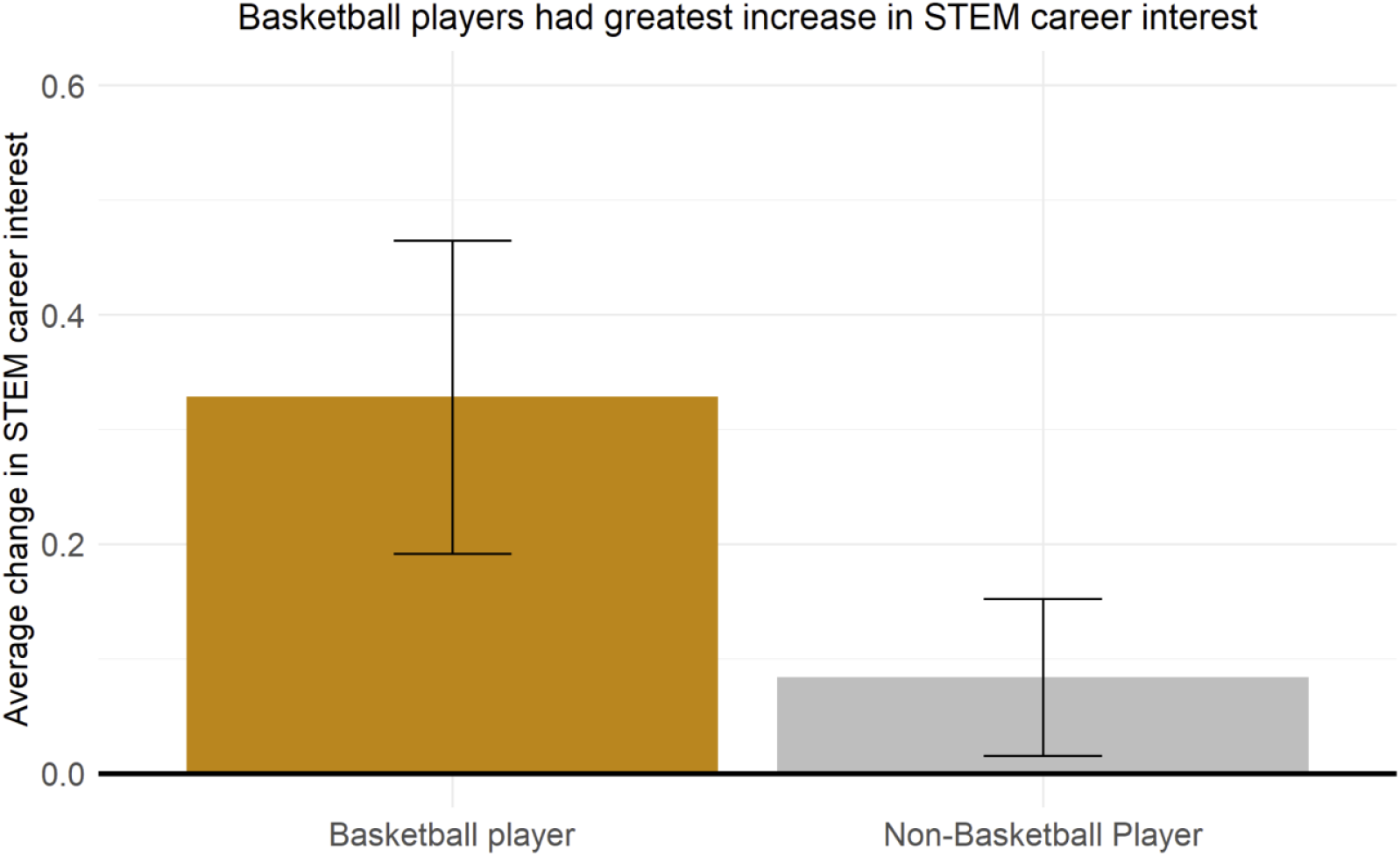
Greatest increase in STEM career interest among basketball players. **Note:** The significant moderation of program effects between basketball players and non-basketball players only applies to Study 1. No moderation analyses were run in Study 2 nor Study 3, due to homogenous sample of basketball players.

The increase in STEM career interest did not significantly differ between students who were already participating in STEM extracurriculars and those who weren’t, *F*(1, 848.67) = 2.40, *p* < .122.

#### STEM enjoyment

Paired t-test results revealed a small but significant increase of STEM enjoyment from prior to participating in the Science of the Slam (*M* = 3.30, *SD* = 0.91) to after (*M* = 3.39, *SD* = 0.88), *M*_*D*_ = 0.09, 95% CI [0.05, 0.13], *t*(784) = 4.16, *p* < .001, *d* = 0.10. This increase in STEM enjoyment did not significantly differ between students already participating in STEM activities and those who weren’t, *F*(1.737.63) = 2.38, *p* =.123, nor between basketball players and non-basketball players, *F*(1.846.95) = 2.52, *p* = .122. However, the increase in STEM enjoyment significantly differed between students who didn’t play sports and those who did, *F*(1.834.14) = 10.09, *p* = .002. STEM enjoyment did not significantly change for students who did not play sports, *t*(274) = 0.16, *p* = .869, but STEM enjoyment significantly increased among students who played sports, *M*_*D*_ = 0.12, 95% CI [0.07, 0.18], *t*(452) = 4.21, *p* < .001, *d* = 0.14.

#### STEM self-efficacy

A Wilcoxon signed rank test revealed no significant change in STEM self-efficacy from before the Science of the Slam (*M* = 3.38, *SD* = 1.01), to after the Science of the Slam (*M* = 3.40, *SD* = 0.95), *Z* = -0.57, *p* = .573.

### Effects on Sports Science Motivation

Sports science interest significantly increased from before the Science of the Slam (*M* = 3.28, *SD* = 1.17), to after the Science of the Slam (*M* = 3.41, *SD* = 1.14), *Z* = 4.14, *p* < .001. The rank-biserial correlation was calculated as 0.15, indicating a small effect size.

There was also a significant increase in how much students valued sports science from before the Science of the Slam (*M* = 3.23, *SD* = 1.01), to after the Science of the Slam (*M* = 3.62, *SD* = 0.96), *Z* = 11.12, *p* < .001. The rank-biserial correlation was calculated as 0.37, indicating a medium effect size.

### Effects on Sport Motivation

There was a small but significant increase in sport enjoyment from before participation in Science of the Slam (*M* = 3.98, *SD* = 0.95) to after (*M* = 4.03, *SD* = 0.96), *M*_*D*_ = 0.04., 95% CI [0.00, 0.08], *t*(793) = 2.01, *p* = .045. This increased sports enjoyment did not significantly differ between students who already were participating in STEM extracurriculars and those who were not, *F*(1, 721.90) = 0.05, *p* = .829, between students who played sports and those who did not, *F*(1, 842.86) = 0.00, .996, nor between basketball players and non-basketball players, *F*(1, 842.23) = 2.33, *p* = .127.

There was no significant change in sports self-efficacy from before the Science of the Slam (*M* = 3.47, *SD* = 1.12), to after the Science of the Slam (*M* = 3.51, *SD* = 1.09), *Z* = -1.87, *p* = .062.

#### Investigating Cross-Domain Motivation Spread

We investigated whether the increase in STEM career interest might be explained by sport interest translating to STEM interest. To do so, we ran a mixed-effects model where STEM career interest was predicted by sports science interest, time (pre vs. post measures), and the interaction between sports science interest and time. There was a significant interaction between sports science interest and time in predicting STEM career interest, *β* = −0.17, 95% CI [−0.21, −0.12], *t* = −6.94. Investigating this interaction revealed that the correlation between sports science interest and STEM career interest was significantly stronger after participation in the Science of the Slam, *r* = .58, 95% CI [.55, .62], *t*(1261) = 25.46, *p* < .001, than before *r* = .39, 95% CI [.35, .43], *t*(1723) = 17.37, *p* < .001 (see Figure 4).

**Figure 4.**
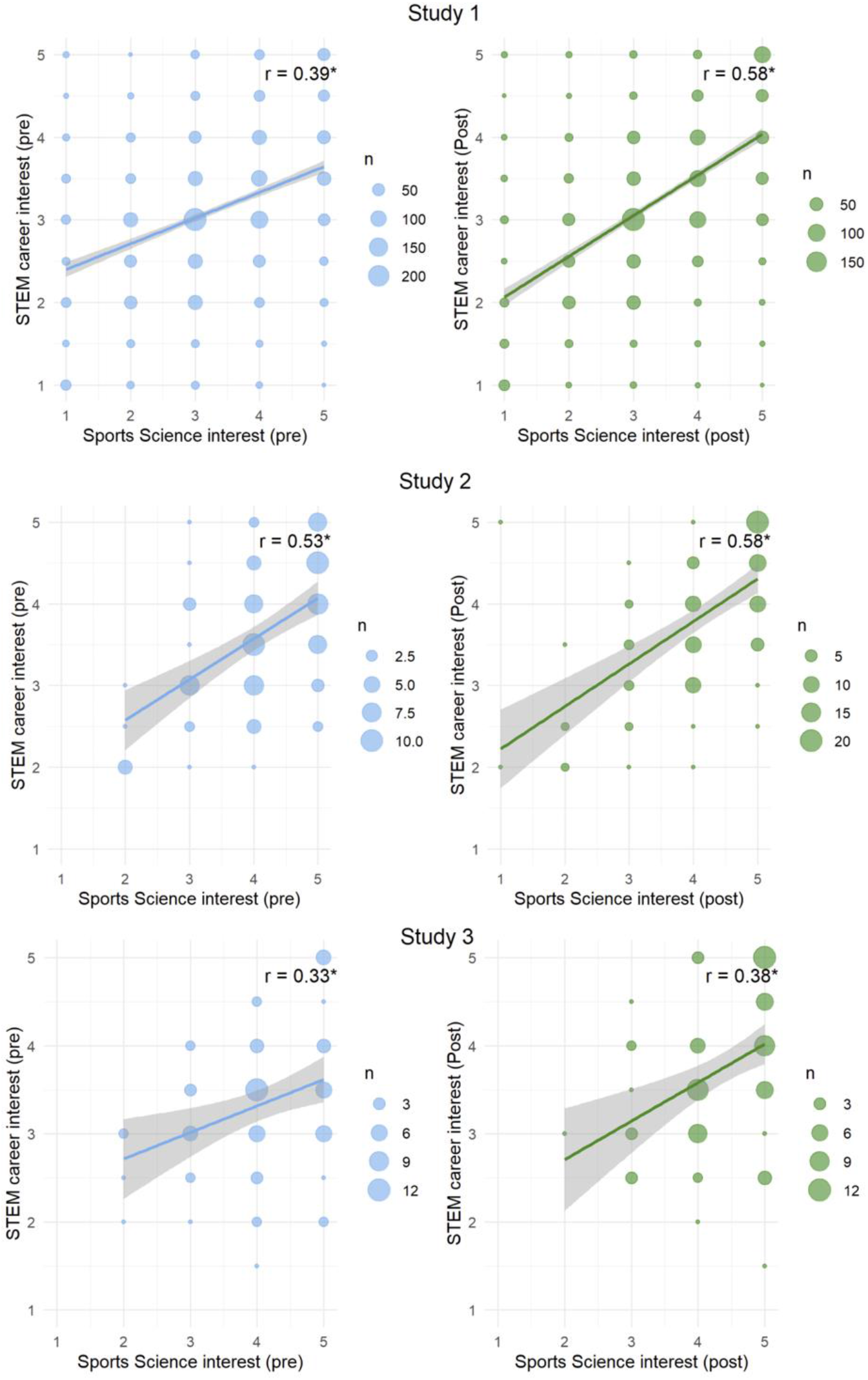
Sports Science interest and STEM career interest correlation (pre vs. post). **Note:** * denotes p < 0.05.

## Discussion

Despite only lasting an hour, the Science of the Slam program increased participants’ enjoyment of STEM activities and furthermore, their interest in pursuing a STEM career.

Notably, the increased interest in pursuing a STEM career was most pronounced for students who played basketball and the increase in both participants’ enjoyment and interest in STEM was driven by student athletes. This suggests that this program was working as intended–its benefits were targeted towards sport-interested youth. Additionally, exploratory mixed-effect model results suggest that the participants’ increased interest in STEM careers did not emerge as an independent interest, but might have been rooted in their pre-existing sports science interest.

Although the Science of the Slam had measurable impacts on participants’ enjoyment and interest in STEM, there was no significant change in participants’ self-efficacy, or their self-assessment of their performance in math and science. This is perhaps not surprising given the relatively brief duration of the Science of the Slam. It would be challenging for a single, one-hour event to compete with years of math and science courses in shaping participants’ perceptions of their math and science capabilities.

In addition to investigating how the Science of the Slam impacted STEM motivation, secondary analyses were conducted to explore any changes to participants’ motivation in sports science and sports. For sports science, participants valued and had greater interest in sports science following the Science of the Slam. As for sports, there was statistically significant but very small increase in sport enjoyment and no change in sport self-efficacy.

Taken together, these findings suggest that the Science of the Slam was effective in its primary aim to enhance interest in STEM among sport-interested youth.

### Study 2

In Study 2 we sought to replicate and extend the results of Study 1 on STEM motivation, by administering a similarly designed but more time-intensive, basketball-based STEM education intervention. Unlike the brief Science of the Slam, this intervention consisted of a one-day clinic. During this four-hour basketball clinic, participants were introduced to the tools and techniques used by sport-scientists to evaluate athletic performance. Mimicking a typical basketball clinic format, youth athletes rotated through stations to participate in various basketball drills and sport-science measurement activities (Loya et al., 2023b). Student participants were recruited through word-of-mouth via local basketball programs, resulting in a primarily basketball-oriented sample of participants.

## Methods

### Participants

A total of 163 students recruited from the Racine and Chicago area participated in a 1-day clinic in each city (*M*_*Age*_ = 11.40 years, *SD*_*Age*_ = 2.38 years). 131 students completed the pre-survey, 152 students completed the post-survey, and 120 students completed both surveys. Students ranged from 2nd grade to 12th grade (see Table 1).

### Materials

The same evaluation materials used in Study 1 were used in Study 2. These included pre and post measures of STEM career interest, STEM enjoyment, STEM self-efficacy, sports science interest, value of sports science, sport enjoyment and sport self-efficacy.

### Procedure

Students completed measures immediately before and after participating in the clinic. Each of the six sport science stations measured a specific basketball attribute measured in the NBA Rookie Combine (speed, quickness, strength, vertical jump height, agility and shooting). For details on a similar program, please see (Loya et al., 2023b). For example, similar to the Science of the Slam in study 1, at the vertical jump height station, participants used a custom-built jump-plate to measure how high they can jump. Using a combination of concepts from computer science and physics, athletes observed how “time in flight” is converted to “height” in real-time. The students logged their data from each station into a provided worksheet for later analysis. Each combine station concluded with a discussion on how to improve performance in a specific athletic domain. For example, for the jump height measurement, the discussion centered around the role vertical jump height plays in basketball, and examples of exercises you can complete at home to improve your jumping ability.

### Data analysis

We tested for program effects by conducting paired t-tests (for composite scores) and Wilcoxon signed ranks tests (for single-item measures) on pre and post survey responses. Initial group differences and moderation by group status are not reported here as the sample was relatively homogeneous and group sizes were not large enough for appropriate comparison (e.g. only 11 students participated in STEM extracurriculars, only 34 students were not basketball players, etc.). To ensure data quality, the same attention check was used as in Study 1. This resulted in the removal of 38 students for a new total of 125 students.

## Results

### Effects on STEM Motivation

There was a significant increase in STEM career interest from before participation in clinic (*M* = 3.62, *SD* = 0.84) to after (*M* = 3.90, *SD* = 0.86), *M*_*D*_ = 0.23, 95% CI [0.07, 0.39], *t*(87) = 2.91, *p* = .005, *d* = 0.27 (see Figure 2). However, there was no significant increase in STEM enjoyment from prior to participating in the clinic (*M* = 3.81, *SD* = 0.76) to after (*M* = 3.92, *SD* = 0.82), *t*(86) = 1.58, *p* = .117, *d* = 0.15. Additionally, there was no significant change in STEM self-efficacy from before the intervention (*M* = 3.92, *SD* = 0.90), to after the intervention (*M* = 3.98, *SD* = 0.84), *Z* = -0.45, *p* = .627.

### Effects on Sports Science Motivation

Sports science interest did not significantly change from before the intervention (*M* = 4.10, *SD* = 0.90), to after the intervention (*M* = 4.19, *SD* = 0.96), *Z* = 0.75, *p* = .484. However, participants’ value of sports science did increase from before the intervention (*M* = 3.88, *SD* = 0.86), to after the intervention (*M* = 4.27, *SD* = 0.82), *Z* = 3.02, *p* = .002. The rank-biserial correlation was calculated as 0.33, indicating a medium effect size.

### Effects on Sport Motivation

There was no significant change in sport enjoyment from before participation in clinic (*M* = 4.63, *SD* = 0.55) to after (*M* = 4.69, *SD* = 0.48), *t*(87) = 0.36, *p* = .720. There was also no significant change in sports self-efficacy from before the intervention (*M* = 4.17, *SD* = 0.87), to after the intervention (*M* = 4.32, *SD* = 0.91), *Z* = -0.72, *p* = .533.

### Investigating Cross-Domain Motivation Spread

As in Study 1, we ran a mixed-effects model where STEM career interest was predicted by sports science interest, time (pre vs. post measures), and their interaction. There was no significant interaction between sports science interest and time in predicting STEM career interest, *β* = 0.02, 95% CI [−0.14, 0.17], *t* = 0.20, suggesting that the correlation between sports science interest and STEM career interest did not significantly increase following the clinic. However, there was both a significant correlation between sports science and STEM career interest before the clinic, *r* = .53, 95% CI [. 37, .66], *t*(95) = 6.13, *p* < .001 and after the clinic, *r* = .58, 95% CI [. 44, .69], *t*(102) = 7.18, *p* < .001, and the correlation coefficient was descriptively larger following the clinic (.58 > .53)(see Figure 4).

## Discussion

Consistent with study 1, STEM career interest significantly increased following participation in the basketball-focused STEM clinic. Notably, only STEM career interest increased, there was no effect on STEM enjoyment nor STEM self-efficacy. Furthermore, this increase in STEM career interest was descriptively larger effect size (*d* = 0.27) relative to Study 1 (*d* = 0.16), despite the smaller sample size for the clinic. There are at least two reasons we speculate may explain why this effect on STEM career interest was descriptively stronger among participants in the clinic. First, the day clinic was more intensive than the single-hour Science of the Slam event, lasting 4 hours (versus 1 hour) and consisting of 6 different sports science activities (vs. 1 activity). This larger effect could simply be due to a higher intervention dosage. Second, the sample in Study 2 was predominantly basketball players (76%) and as indicated in the results of Study 1, the effect of the informal STEM intervention on STEM career interest is most pronounced among basketball players.

Also consistent with Study 1, the value of sports science increased among participants after participating in the clinic. However, unlike Study 1, there were no significant changes in any of the sports-related measures (i.e., enjoyment of sports nor self-efficacy in sports). The absence of a significant change in sports enjoyment and sports self-efficacy is likely due to ceiling effects born out from the particularly sports-engaged sample of Study 2. Indeed, the average sports enjoyment rating prior to the clinic was already at 4.63 (out of 5.00). This may indicate that this sample was already maximally engaged in basketball, just as participants in many informal STEM programs are already maximally engaged in STEM programming (Staus et al., 2021).

### Study 3

In Study 3 we administered the most time-intensive basketball-based STEM education intervention as a three-day camp, with each session consisting of six hours of instruction (for a total of 18 hours). The guiding principle of the camp was to showcase basketball’s inherent STEM elements, redefine STEM within a familiar environment, and inspire athletes to pursue careers at the intersection of STEM and sports by equipping students with tools to design their own basketball-centered research experiment. The program was designed to mimic the structure and atmosphere of a traditional basketball camp to appeal to students who have an existing affinity for basketball and who aspire to improve their athletic abilities. The camp incorporated basketball stations for targeted fundamental skill development (shooting, passing, dribbling, etc.), controlled game-like scenarios (3 vs. 3 competitions, 2 vs. 1 advantage situations, defensive rotations, pick and roll techniques, etc.), and full court scrimmages to synthesize and apply all competencies. In addition to the pure basketball content, we integrated STEM lessons, language, and activities into each drill. For example, we used principles from mechanics to adjust sprinting form, we applied basic statistics to evaluate shooting efficiency, and we quantified physical athletic attributes in combine stations using low-fidelity sport science equipment.

Throughout the duration of the camp, students progressed from being passive recipients of basketball and STEM instruction, to becoming active participants who were prepared to question, analyze, and generate new knowledge in ways that enhanced their performance on the basketball court. To accomplish this, each day of the camp was assigned a distinct theme. On the first day, the focus was to show students how the scientific method can be applied to answer questions related to sports performance, and expose them to common experimental tools used in the fields of musculoskeletal, orthopedic, and biomechanics research. The aim of the second day was to introduce students to aspects of the engineering design process, sports analytics methods, experimental design, and data visualization techniques. These lessons served as prerequisites for the basketball-centered research project that the students would design, conduct, and present as part of a conference-style poster session for an audience of local coaches, musculoskeletal research experts, and peers on the last day of the camp.

## Methods

### Participants

A total of 98 students recruited from the Chicago and Racine areas participated in the camp (*M*_*Age*_ = 12.28 years, *SD*_*Age*_ = 1.84 years). 84 students completed the pre-survey, 97 students completed the post-survey, and 83 students completed both surveys. Students ranged from 4th grade to 11th grade (see Table 1).

### Materials

The same materials were used as described in Study 1 and Study 2.

### Procedure

Students completed measures immediately before and after participating in the camp. At the start of each day of the camp, local university, junior college, and high school basketball coaches led the students through a series of dynamic warm-up exercises, which are commonly performed prior to a basketball competition in order to prime the body for subsequent sport-specific movements. Rather than simply directing the student athletes to complete a sequence of movement patterns, coaches identified which muscle groups were targeted by each exercise and explained how each movement corresponded to specific basketball skills. Building upon the warm-up series, students also participated in mini competitions that emphasized how proper technique impacts both linear and lateral movements to fully realize the underpinning biomechanics principles. The introductory portion of each day concluded with remarks from leading experts in both professional basketball and STEM to 1) corroborate the value of the forthcoming learning objectives and 2) serve as role models the youth athletes can identify with. The following descriptions detail the STEM curriculum, omitting the basketball training segments, which were interwoven throughout each day.

#### Day 1 – Scientific Method & Biomechanics

Students completed each of the six NBA Rookie Combine stations (speed, quickness, strength, vertical jump height, agility and shooting) described in Study 2. In addition to recording their own performance results, students were provided with a deeper explanation of the engineering behind each low-fidelity sports science tool. They also brainstormed possible basketball questions that could be answered with their data from the sport science equipment.

#### Day 2 - Engineering & Sports Analytics

Students separated into groups of about four students to form individual research teams, led by one coach who would serve as the advisor for the project. Each group aligned on a research question they wanted to ask and selected the appropriate sports science tool that would help them answer their question. One example of a research question was, “How much of an effect does fatigue have on vertical jump height?” The team measured vertical jump height before and after instructing their study participants to perform an isometric squat hold for two minutes. The team calculated the percent change in vertical jump height, identified the physiological principles underpinning their results, and drew conclusions related to how these results could inform their basketball training. Finally, using examples from the NBA Rookie Combine stations as inspiration, the students decided how they would represent their findings in the form of a table, graph, etc.

#### Day 3 – Poster Preparation & Presentations

Coaches provided student teams with a poster template, on which students filled in their research project title, question, hypothesis, methods, results, physiological principle, conclusions, and future work. Students rehearsed their respective speaking roles with the staff. The camp concluded with a conference-style poster session for invited guests such as local coaches, teachers, and biomechanics researcher as well as an academic award ceremony to celebrate the participants’ commitment and growth within sports and STEM.

### Data analysis

Data analyses and the attention check were the same as in Study 1 and 2, for a new total of 86 students.

## Results

### Effects on STEM Motivation

There was a moderate and significant increase in STEM career interest from before participation in camp (*M* = 3.33, *SD* = 0.79) to after (*M* = 3.73, *SD* = 0.85), *M*_*D*_=0.41,, 95% CI [0.19, 0.63], *t*(70) =3.71, *p <*.0.001, *d* = 0.50 (see Figure 2). However, there was no significant change in STEM enjoyment from prior to participating in the camp (*M* = 3.57, *SD* = 0.79) to after (*M* = 3.69, *SD* = 0.78), *M*_*D*_=0.08, 95% CI [−0.06, 0.21], *t p* =.281, nor in STEM self-efficacy from before the camp (*M* = 3.84, *SD* = 0.82), to after the camp (*M* = 3.90, *SD* = 0.93), *Z* = -0.36, *p* = .737.

### Effects on Sports Science Motivation

Sports science interest significantly increased from before the camp (*M* = 4.05, *SD* = 0.86), to after the camp (*M* = 4.38, *SD* = 0.74), *Z* = 3.03, *p* = .003. The rank-biserial correlation was calculated as 0.34, indicating a medium effect size. The value of sports science significantly increased from before the camp (*M* = 3.71, *SD* = 0.86), to after the camp (*M* = 4.40, *SD* = 0.76), *Z* = 5.08, *p* < .001. The rank-biserial correlation was calculated as 0.54, indicating a medium-to-large effect size.

### Effects on Sport Motivation

There was no significant change in sport enjoyment from before participation in camp (*M* = 4.70, *SD* = 0.48) to after (*M* = 4.77, *SD* = 0.44), *t*(72) =1.72, *p* = .0.90 . There was no significant change in sports self-efficacy from before the camp (*M* = 4.23, *SD* = 0.81), to after the camp (*M* = 4.38, *SD* = 0.80), *Z* = -1.03, *p* = .316.

### Investigating Cross-Domain Motivation Spread

The correlation between sports science and STEM career interest did not appear to significantly increase following the camp, as there was no significant interaction between sports science interest and time in predicting STEM career interest, β = −0.09, 95% CI [−0.36, 0.17], *t* = −0.70 . However, there were significant correlations between sports science interest and STEM career interest both before the clinic, *r* =.33, 95% CI, [.11, 52], *t*(70) = 2.95, *p* =0.004 and after the clinic, *r* =.38, 95% CI, [.18, .55], *t*(18) =3.71, *p* <.001, and the correlation coefficient was descriptively larger following the camp (.38 > .33) (see Figure 4).

## Discussion

Consistent with Study 1 and 2, STEM career interest significantly increased following participation in the camp activity. Like Study 2, only STEM career interest increased among the STEM motivation variables, with no significant change in enjoyment of STEM nor STEM self-efficacy. Most importantly there appear to be dosage impacts, because as the most time-intensive intervention across the three studies, the 3-day camp lead to the descriptively largest effect size on STEM career interest (*d* = 0.50) compared to the effects of the 1-hour intervention in Study 1 (*d* = 0.14) and the effects of the 4-hour clinic in Study 2 (*d* = 0.27), despite Study 3 having the smallest sample size (N = 86 participants). Likewise, the results of Study 3 were consistent with Study 2 as the value of sports science significantly increased and no significant changes were observed in sports enjoyment nor sports self-efficacy.

### General Discussion and Conclusions

Our novel use of sports as a venue for STEM education approach was designed to not ask youth to conform to existing STEM identities, but instead to authentically embed STEM practices in a domain within which they already possess cultural capital. Sports provide a transformative venue, which reimagines what STEM actually means to these student-athletes, offering an entryway for youth who are not attracted to traditional STEM programs due to a mismatch of cultural capital. Importantly, sports are already culturally relevant to millions of youth and offer an accessible platform to integrate STEM education into activities these youth already care deeply about and feel ownership of. Our finding in Study 1 that students who already reported participating in STEM programming had a statistically significant higher STEM interest (*M* = 3.50, *SD* = 0.93) relative to the general student population (*M* = 3.10, *SD* = 0.93) speaks to the self-selected nature of traditional “STEM-Intensive” programming (Archer et al., 2015; Staus et al., 2021) and the need for more broad-based approaches for participant recruitment.

The growth we observed in STEM career interest was statistically significant across all three studies, with an increased effect size as the duration of the program increased. Interestingly, the primary ceiling effects that we observed in our STEM intervention was not among STEM motivational variables but among sports-related indices, as our samples in Study 2 and 3 were heavily recruited from sport-active populations. Similar to traditional informal STEM education programs, we recruited a biased participant pool; however, it was biased based on a pre-existing interest in sports and a lack of STEM interest. Thus, we were able to demonstrate an effective intervention for enhancing STEM interest among those who would most benefit and did so by creating connections between existing interests in sports and transmuting them into an interest in sport science and, more broadly, STEM careers generally.

Anecdotally, student identity and ownership of sports was very apparent among participants in our STEM interventions. At the start of the Science of the Slam program (Study 1), the student group are asked “what is a hypothesis?” Students often remained quiet and then at the prompting of a teacher or coach, one would tentatively raise their hands and make their own hypothesis: “An educated guess?” In contrast, after introduction to the scientific method through the framing of a slam dunk contest and measurements of each subject’s vertical jump, when asked “What is your hypothesis, who is going to win and why?” There was typically a wild clamor of raised hands and suggestions. Similarly, in the camp (Study 3), when the campers were first told on day 1 that they would be making a poster, there were a few murmurs of discontent, “I’m here to play ball, not do school.” However, by the end of the program at the poster session, the same camper very proudly shared their findings that the loading period in a countermovement jump was an important contributor to vertical jump height and declared that “This’ll have me dunking by next year, latest.” When considering that this student is a 4’6” middle schooler and the audience was a mix of engineering professors, peers, and coaches, it really highlights the confidence that these students felt within basketball and, by the end of the program, STEM within sports. It is difficult to imagine any middle school student holding court with a crowd of STEM professionals in such a confident manner about a traditional “STEM-Intensive” research project such as robotics. We posit that the difference is that students feel ownership of their own sports performance and knowledge.

As promising as our results are, there are notable limitations of these studies. First, the samples used in these studies were very specific and targeted samples—youth recruited from the Chicago and Racine area that were generally sports-engaged participants. Although we consider the specificity of the samples to be a strength as they capture a particularly STEM-uninvolved population, results from these studies should not be generalized to a broader population without appropriate testing in more diverse samples. Likewise, this STEM intervention was tailored to this population by focusing on a sport popular in this population (e.g., basketball). It is highly unlikely and unadvised for adopters of this STEM intervention to administer it in populations where interest in basketball is not particularly strong. Indeed, we recommend and urge researchers to consider culturally aligning STEM interventions to the particular interests of their targeted populations and encourage building upon this current work by implementing STEM interventions that utilize other domains often overlooked in traditional STEM informal education programs (e.g., dance, music, cooking, etc.).

Overall, what likely makes this current STEM intervention effective is its ability to meet students where they are, both physically and culturally. Sports-based STEM programs often happen in gyms, schools, and community centers—places where youth arguably already feel a sense of belonging. Such programs can help ameliorate barriers like cost and transportation, which limit access to traditional STEM initiatives. Ultimately, by expanding the domains of informal STEM programs, we are able to expand the reach of STEM exposure and potentially redefine who can thrive in science and technology careers.

## Declarations

### Availability of data and materials

The datasets used and/or analyzed during the current work are available from the corresponding author on reasonable request.

### Authors’ contributions

EH: Conceptualization, Data curation, Formal analysis, Writing - Original Draft Preparation, Writing - Review & Editing; JD: Conceptualization, Investigation, Writing - Original Draft Preparation, Review & Editing; AL: Conceptualization, Investigation, Writing – Original Draft Preparation, Review & Editing. All authors read and approved the final manuscript.

## Acknowledgements

We would like to thank 4^th^ Family Inc. for their long-term collaboration on sports for STEM education programming and on this project. Specifically, we want to acknowledge 4^th^ Family co-founders, John Scott and Jahkeen Hoke, as well as Jesse Smith Jr., Jamel Edwards, Phillip Smith, John Minogue, Leigh Klein, Nicole Conley, Kennedy Clark, Megan Lee, and all of our other volunteers for assistance running the program. We would also like to acknowledge Caroline Dettman and Michelle Castle from Have Her Back Consulting for their assistance. Finally, we would like to Jim Ladwig, Alan VanderMOlen, and Kristen Beglinger from SC Johnson for their support for this program.

